# Designing and implementing programmable depletion in sequencing libraries with DASHit

**DOI:** 10.1101/2020.01.12.891176

**Authors:** David Dynerman, Amy Lyden, Jenai Quan, Saharai Caldera, Aaron McGeever, Boris Dimitrov, Ryan King, Giana Cirolia, Michelle Tan, Rene Sit, Maarten van den Berge, Huib A. M. Kerstjens, Alen Faiz, Stephanie Christenson, Charles Langelier, Joe DeRisi, Emily Crawford

**Affiliations:** Chan Zuckerberg Biohub, San Francisco, California, USA; Division of Infectious Diseases, University of California San Francisco, San Francisco, California, USA; Department of Biochemistry and Biophysics, University of California San Francisco, San Francisco, California, USA; Department of Microbiology and Immunology, University of California San Francisco, San Francisco, California, USA; Chan Zuckerberg Initiative, Redwood City, CA; Department of Pulmonary Diseases, University of Groningen, Groningen, The Netherlands; University of Technology Sydney, Respiratory Bioinformatics and Molecular Biology (RBMB), School of Life Sciences, Sydney, Australia; Division of Pulmonary, Critical Care, Allergy and Sleep Medicine, University of California San Francisco, San Francisco, California, USA

## Abstract

Since Next-Generation Sequencing produces reads uniformly subsampled from an input library, highly abundant sequences may mask interesting low abundance sequences. The DASH (Depleting Abundant Sequences by Hybridization) technique takes advantage of the programmability of CRISPR/Cas9 to deplete unwanted high-abundance sequences. Because desired depletion targets vary by sample type, here we describe DASHit, software that outputs an optimal DASH target set given a sequencing dataset, an updated DASH protocol, and show depletion results with DASHit-designed targets for three different species.

## Main Text

Next Generation Sequencing (NGS) of RNA libraries produces reads from an unbiased subsample of the input, so highly abundant sequences dominate NGS output and potentially prevent detection of interesting low abundance sequences. Previously, we described DASH^1^ (Depletion of Abundant Sequences by Hybridization), a CRISPR/Cas9 technique that cleaves user-defined sequences targeted by guide RNAs (gRNAs),^2^ rendering them unsuitable for binding to a sequencing flow cell (**Figure 1a**). By using DASH to reduce known high-abundance background sequences, greater sequencing depth is achieved for sequences of interest, enabling detection of, for example, low-abundance pathogens, anti-microbial resistance genes, and transcripts with low expression levels. Importantly, in our prior work, we demonstrated preservation of the relative abundances of human transcript sequences, confirming that DASH can be used for gene expression studies.^1^ Previously we demonstrated that using DASH can amplify low-abundance pathogens in human metagenomic cerebrospinal fluid by 3- to 10-fold.^1^ Other groups have applied DASH to reveal low abundance pathogens in clinical settings,^3–6^ and developed similar techniques to deplete known mitochondrial sequences and thus focus sequencing depth on nuclear sequences of interest in chromatin studies.^7,8^

**Figure 1:**
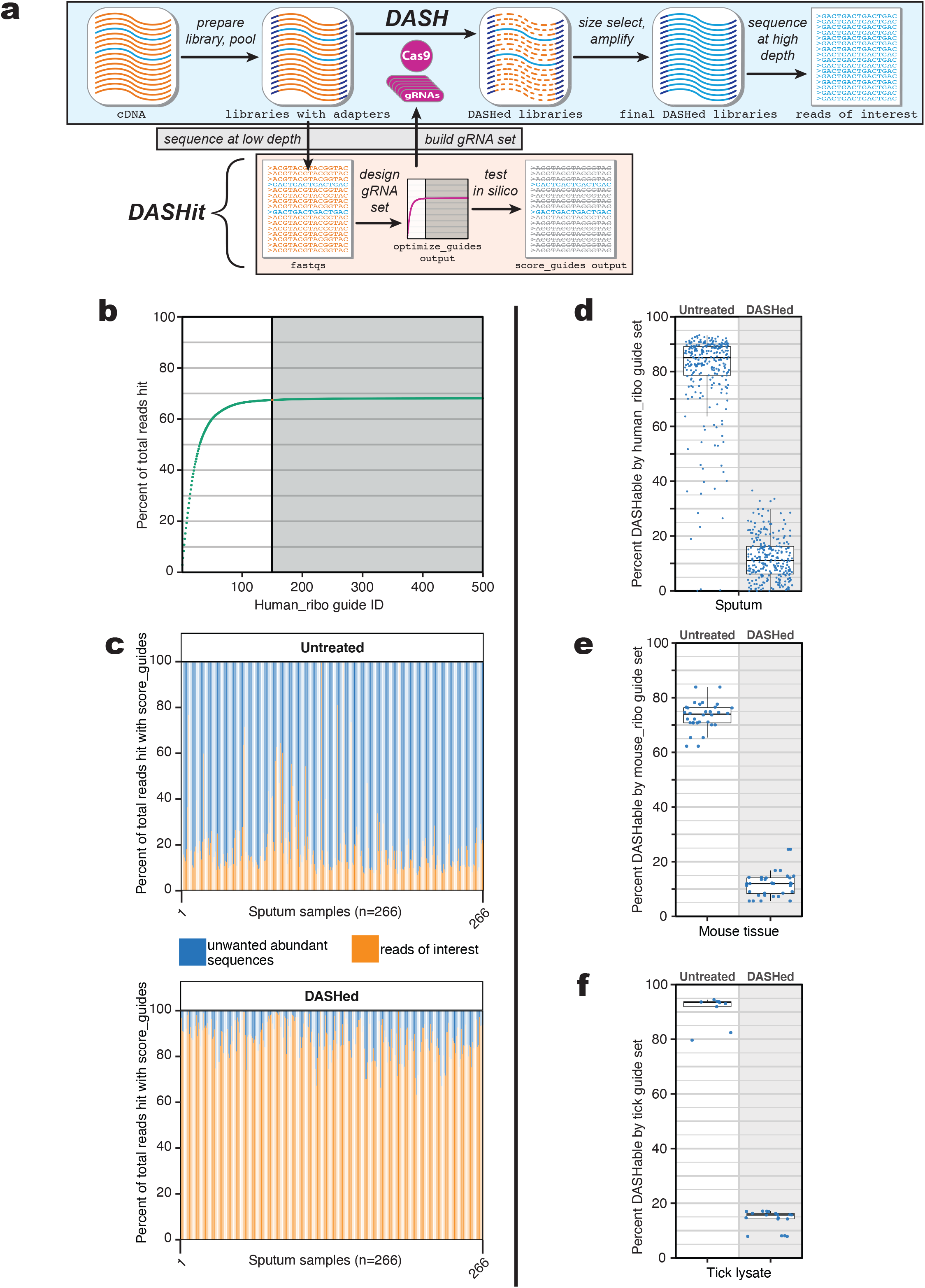
**a.** DASH protocol schematic, including sequencing at a low depth for DASH guide design and pooling prior to DASH. **b.** Output from running DASHit on human sputum samples to target ribosomal RNA. The “elbow plot” illustrates diminishing depletion returns with increasing guides. 150 guides hit 67.4% of total reads and 500 guides hit 68.1% of total reads. **c.** Percent of total reads in untreated and DASHed sputum samples divided into two groups: unwanted abundant reads that are hit by `human_ribo` guide set as measure by `score_guides`, and all others (reads of interest). **d.** Boxplot illustrating `score_guides` results in both untreated and DASHed sputum samples with the `human_ribo` guide set. Each dot is a sample. **e**,**f.** Boxplots illustrating `score_guides` results in both untreated and DASHed samples with the appropriate guide set. Each dot is a sample.

These depletion methods all require time-consuming manual design of the CRISPR/Cas9 gRNAs that will target highly abundant sequences, which vary by library prep method and sample type. To remove this limitation and broaden the applicability of the technique, we developed DASHit, an automated guide design procedure, together with a collection of open source software tools to build DASH gRNA sets and predict the success of a DASH run, as well as an improved DASH wet lab protocol. These improvements allow researchers to easily apply NGS to any study where uninformative high abundance sequences mask biologically interesting questions. Importantly, DASHit enables ribosomal RNA depletion in non-model organisms for which commercial depletion kits are not available.

DASH is most effective when a small number of guides target a large proportion of highly abundant reads - for example, in RNA-seq libraries where ribosomal reads dominate. In contrast, DASH is less effective on DNA-seq libraries, where typically no particular sequence type dominates and an intractable number of guides would be required for effective depletion. DASHit designs gRNA sets by processing an initial low-depth sequencing run that contains the over-abundant sequences the user would like to deplete and automatically choosing a small set of gRNAs that targets them. The user provides an adaptor-trimmed .fasta file of initial sequencing reads `input` and specifies an integer `num_guides > 0`. Then, DASHit finds a set of `num_guides` gRNAs that hit the largest number of reads in `input`. Finding such a set is an instance of the “Maximum Coverage Problem,” a well-known problem studied in computer science that cannot be efficiently solved in general. Instead, we adopt the greedy approach, which is known to be a good approximation to the optimal solution,^9^ and build a set of `num_guides` gRNAs by repeatedly choosing a gRNA that hits the largest number of reads in `input`, removing the reads hit from consideration, and then repeating. In this fashion, we produce a guide set that targets the most abundant reads in `input`. Additionally, DASHit outputs a running tally of the number of reads hit by guides as each additional gRNA is added to the set, providing a visual tool (**Figure 1b**) for evaluating the utility of DASH and selecting an optimal size for the gRNA set.

During guide design DASHit allows the user to filter gRNAs by specifying off-target regions (guides matching off-targets will not be selected) and on-target regions (only guides in on-target regions will be considered) in the form of .fasta files, which can be prepared from reference sequences using BEDTools.^10^ DASHit can further filter guides for quality, where gRNAs with polymeric repeats, extreme GC content and hairpins are excluded.

After designing guides, DASHit can score the resulting set of gRNAs. The program `score_guides` takes as input an existing set of gRNAs and a sequence file and reports the number of reads that are targeted by at least one gRNA. In addition to verifying that the selected guides hit the targets of interest in the initial sequencing run, `score_guides` is useful for estimating how well an existing guide set will perform against a new sample.

The DASHit software, including source code, is freely available at http://dashit.czbiohub.org under the open-source Apache 2.0 license. This website also contains examples of guide design and helper scripts for batch processing. The Online Methods section contains a complete description of all the algorithms in DASHit. We also present an updated lab protocol, improved from our earlier method^1^ in several ways: (1) it is applicable to both Nextera- and TruSeq-style library prep methods; (2) it is applied after completion of library prep, so already-sequenced libraries can be DASHed and re-sequenced; (3) it can be applied to pooled libraries; and (4) clean-up no longer requires phenol/chloroform, improving automation compatibility and reducing hazardous waste. Our complete improved protocols are available on protocols.io^11,12^.

To validate the guide design procedure described above, we used DASHit to design three different guide sets for samples from human, mouse, and tick. We then performed DASH and measured each set’s effectiveness using `score_guides`.

First we considered a set of human sputum samples from a study designed to evaluate both host transcriptome and pathogen presence. An initial low-depth RNA sequencing dataset contained a high fraction of reads mapping to human nuclear ribosomal RNA (average 78%), and a smaller fraction to human mitochondrial ribosomal RNA (average 4%) (**Figure S1**). We randomly subsampled 100,000 reads each from 41 sputum libraries and used DASHit to generate gRNAs targeting these reads. To avoid the possibility of hitting transcripts of interest, we used all human ribosomal RNA sequences as an on-target input file for DASHit, and the human genome with those sequences excluded as an off-target input file (see Online Methods). After DASHit designed 500 guides, we examined the “elbow plot” produced (**Figure 1b**) and found that beyond 150 gRNAs the benefit of including additional guides dropped off. Thus we chose the 150 guides that hit the largest number of reads in the initial RNA-seq run and named this set `human_ribo`. To make the guide RNAs, we purchased DNA templates for the CRISPR RNA (crRNA) corresponding to each, pooled them, and performed *in vitro* transcription as described in Online Methods.

Next, a pooled library of 266 sputum samples was subjected to DASH using `human_ribo` gRNAs as described in Online Methods. The untreated and DASHed libraries were each sequenced on an Illumina MiSeq, with a median 42,270 PE150 read pairs collected per sample. We ran `score_guides` with `human_ribo` on both the untreated and DASHed libraries for each sputum sample and observed a median 7.7-fold (range 2.5-76.7) decrease in the presence of these target sites following DASH (**Figures 1c** and **1d**). Using bowtie2 we confirmed that the reads depleted were ribosomal (**Figure S1**). For 22 of these samples, a respiratory virus was present at 1 read per 10,000 read pairs or greater; viral reads per 10,000 reads increased a median of 8.6-fold in the DASHed samples (range 1.6-23.4, plus 4 samples with less than 1 per 10,000 reads in the untreated control) (**Table S1**). We conclude that our DASHit designed guides allow for more efficient detection of viral pathogens in metagenomic samples.

Next, using the same method described above, we used DASHit to design a set of 96 mouse ribosomal DASH gRNAs based on 16 tissue samples (stomach, small intestine, large intestine, and cecum) from a transcriptomic study of female BALB/c mice (**Figure S2**). DASH was applied as described in Online Methods, and `score_guides` showed that the targeted sequences decreased from an average of 73% in the original samples to 12% in the DASHed samples (**Figure 1e**), freeing that sequencing space for more biologically interesting portions of the transcriptome. Mouse tissue libraries were kindly provided by Drs. Suzanne Noble and Pauline Basso at UCSF.

Finally, to test how DASHit might perform in a non-model organism, we designed a tick gRNA set based on 19 tick RNA-Seq libraries kindly provided by Dr. Seemay Chou at UCSF. Nine of our RNA-Seq libraries were prepared in various mixtures from a total of 48 individual ticks of the genus *Dermacentor*, various species. The other ten represent individual *Ixodes pacificus* ticks. Because the tick genome has only limited annotations we did not apply on-target or off-target filtering but did manually remove a single candidate gRNA that mapped to the pathogen *Borellia bergdorferi* genome from the final set of 191 gRNAs (**Figure S3**). DASH was applied to the nine *Dermacentor* samples as described above, and `score_guides` showed that the targeted sequences decreased from an average of 91% to 14.1% (**Figure 1g**). This frees up sequencing depth to detect lower-abundance sequences such as pathogens. Additionally, to illustrate the limited ability of DASH to deplete unwanted sequences in DNA-Seq libraries (which don’t have small numbers of highly abundant sequences like RNA-Seq libraries do), we also ran DASHit on a DNA-Seq library prepared from a single *Ixodes scapularis* tick, and found that a set of 500 gRNAs would deplete only 13.9% of reads (**Figure S3**).

DASH has proven useful in several studies^3–6^ since its initial publication, and other groups have developed similar strategies,^7,8^ demonstrating the utility of removing uninformative high-abundance reads in a number of different types of studies. This utility will only increase with the adoption of small sequencers like Illumina’s iSeq, which otherwise produce too little sequencing output for many RNA-Seq studies. The automated guide design approach implemented in DASHit, together with the streamlined wet lab protocol we describe, makes DASH substantially more accessible for researchers applying NGS to studies where uninformative high abundance sequences mask biologically interesting questions.

## Online Methods

Inspired by the UNIX philosophy of chaining together small programs that perform one task, DASHit is made up of several tools: first, the input is preprocessed by `crispr_sites`, which scans input NGS sequencing reads, formatted as a FASTA-file, and outputs all Cas9 target sequences in the input, along with which reads contained each target sequence. Cas9 targets are regions of DNA where the Cas9 gRNA complex can bind. They are 20 nucleotides long and must be directly next to a protospacer adjacent motif (PAM): either NGG on the 3’ end of the target, or CCN on the 5’ end. We think of the resulting output as a candidate list of all possible guide RNAs that would bind to the sequences provided in the input. `crispr_sites` makes use of low-level optimizations possible in C++11, for example using a bitcode encoding to store binding sites, to achieve fast performance: finding all Cas9 binding sites in the 3.2 gigabase human genome takes approximately 2 minutes on a laptop.

The output of `crispr_sites`, together with a requested number of gRNAs `num_guides`, is provided to `optimize_guides`, which finds a set of `num_guides` gRNAs that covers the largest number of reads in the input. This is an instance of the “Maximum Coverage Problem”, a well-known intractable optimization problem. Fortunately, it is also known that a greedy algorithm provides a good approximate solution to this problem: the set chosen by the greedy algorithm will, in the worst case, cover 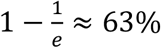 of what is covered by the optimal set;^9^ moreover, this is the best possible approximation ratio for this problem. Our greedy implementation proceeds as follows:

1. Mark all reads in the input as not hit and initialize an empty list of gRNAs `guides`.
2. Pick a gRNA in the input that hits the largest number of not-hit reads, add that gRNA to `guides`, and mark all the reads hit by this gRNA as hit.
3. Repeat Step 2 until `guides` contains `num_guides` gRNAs.

`optimize_guides` outputs `guides`, ordered by the number of input reads hit, together with a running total of the percentage of input reads hit. This running total can be plotted (**Figure 1b**) to estimate how many gRNAs to use before diminishing returns, in terms of the number of input reads hit, sets in.

Prior to running `optimize_guides`, the output of `crispr_sites`, containing all gRNAs found in the input, can be run through `dashit_filter` to optionally remove candidate gRNAs that:

1. have poor structure, meaning:

a. GC content: too few or too many G’s and C’s (default 5 <= #GC <= 15),
b. Homopolymer: a single nucleotide repeat (default 5 < #repeat),
c. Dinucleotide repeat: a repeating pair of nucleotides (default 3 < #repeat), and,
d. Hairpins: a DNA stem-loop (default 5 <= stem length, 3 <= loop length).
2. match a list of off-target gRNAs, or,
3. do not match a list of on-target gRNAs.

off-target and on-target lists of gRNAs are created by processing the desired off-target/on-target sequence files in FASTA format through `crispr_sites`. To extract off-target/on-target gRNAs from, for example, annotated reference sequences, the input FASTA files can be masked with BEDTools^10^ prior to processing with `crispr_sites`.

Finally, the `score_guides` program takes as input a gRNA set produced by `optimize_guides` and a file containing sequencing reads, and reports how many reads are hit by guides in the provided guide set. `score_guides` is primarily useful to estimate how effective guides designed against one sample will be at DASHing a related one.

Next we describe in detail our guide design and validation process using DASHit for the mouse samples. The guide design and DASH experiments for the human and tick samples were processed similarly.

We examined the sequences in FASTQ format from a low-depth initial paired-end 150bp Illumina MiSeq run (greater than 100,000 reads) of mouse RNA-seq libraries created from 16 tissue samples (stomach, small intestine, large intestine, and cecum) from female BALB/c mice. We recommend using 150bp reads to increase the chances of identifying a cut site on a given DNA molecule. We first filtered the FASTQ files for quality using PriceSeqFilter^14^ such that 90% of nucleotides in a read pair must be called (flag `-rnf 90`) and 85% of nucleotides in a read pair must have a 98% probability or higher of being correct (flag `-rqf 85 0.98`). We then converted the sequences to FASTA format and randomly subsampled 100k reads from each of 16 mouse libraries using seqtk^13^, and trimmed TruSeq adaptor sequences using cutadapt.^15^ Trimming adaptor sequences is particularly important, since a DASH guide set targeting this region would destroy the adaptor regions and render a DNA library unsequenceable. After this preprocessing, we ran `crispr_sites` with the `-r` to identify all possible gRNA sites in the input, together with which reads each gRNA appears in.

Before finding an optimal gRNA set, we decided to only include on-target gRNAs that matched rRNA regions in the Genome Reference Consortium^16^ GRCm38/mm10 mouse reference genome, obtained from the UCSC Genome Browser^17^, and remove off-target gRNAs that matched non-rRNA mouse sequences, as well as gRNAs that were present in the *Candida albicans* SC5314 genome^19^ as we were interested in the *C. albicans* transcriptome *in vivo*. We created a bed file indicating all mm10 rRNA genome annotations and used BEDTools to extract these sequences into an on-target FASTA file. We created an off-target FASTA file by using BEDTools to extract the non-rRNA sequences from the mm10 genome and concatenated that with the *C. albicans* genome. Next we ran `crispr_sites` on both these FASTA files to produce a list of on-target and off-target gRNAs. Finally, we performed filtering of our original gRNA list by running `dashit_filter` and providing the on-target and off-target gRNAs. The resulting filtered list of gRNAs represents all those found in our original low-depth sequencing run, except: those not matching the on-target gRNA list, or those matching the off-target list, or those that failed the structure quality tests (structure quality filtering settings were left at their default values). Next, we provided the filtered list of gRNAs to `optimize_guides` to pick the top 500 gRNAs. Examining the elbow plot (**Figure S2**), we chose the top 96 gRNAs based on elbow plot.

A similar process was followed for the design of our human and tick gRNA sets. The human set included filtering to only include on-target guides hitting human rRNA regions of the hg38 human genome as annotated by the UCSC Genome Browser^18^ and to exclude guides with off-target activity on human non-rRNA regions of the hg38 human genome. The tick gRNA set did not include any filtering for on or off target activity. The DASHit software site http://dashit.czbiohub.org hosts more detailed documentation, including wrapper scripts that perform the steps above in one go.

After the guide design process above, we ordered crRNA DNA templates and tracrRNA DNA templates of our Cas9 gRNA sequences plus a 18bp T7 promoter from Integrated DNA Technologies (IDT). Complementary 18bp T7 promoter DNA sequences also from IDT were annealed to the 18bp T7 promoter on the gRNA DNA templates. All crRNAs for a given pool (mouse, human, or tick) were combined in equimolar concentrations and then diluted to 40ng/μL by Nanodrop. For transcription the following were combined: 100μL of 100ug/μL T7 RNA polymerase (prepared in house), 100μL 40ng/μL crRNA template pool or tracrRNA template, 120μL 10X T7 buffer (400 mM Tris pH 7.9, 200 mM MgCl2, 50 mM DTT, 20 mM spermidine), 300μL 25mM each NTPs from Thermo Fisher Scientific, and 380μL of nuclease-free water. Transcription took place at 37°C for 2 hours, and then gRNAs were purified using ethanol precipitation and SPRI bead clean up. crRNA and tracrRNA were diluted to 80μM by concentration from Life Technologies’ Qubit HS RNA kit and annealed together for a final concentration of 40μM. Single use aliquots were stored at −80°C. It is also possible to directly order synthetic gRNAs based on the output of DASHit.

After we transcribed our designed gRNAs, we performed DASH experiments with our guides. Libraries were prepared using NEB Next Ultra II RNA Library Prep Kit for Illumina. Pooled libraries were diluted to a final concentration of 1.4 nM DNA (or 0.42 ng/µL for a library with average fragment size of 450 bp) and incubated with a final concentration of 5 µM *Streptococcus pyogenes* Cas9 (recombinantly expressed and purified from *E. coli* in house, which is described in detail in our previous work^1^) and 10 µM gRNAs in a buffer containing 50mM Tris pH 8.0, 100mM NaCl, 10mM MgCl_2_ and 1mM TCEP at 37°C for 30 minutes. A Zymo DNA Clean and Concentrate column clean was performed, and then the library was DASHed again with the same conditions for 90 minutes. The reaction was quenched with Proteinase K from NEB, cleaned up with SPRI beads at a bead:sample ratio of 1:0.9 to remove DASH-digested fragments, amplified using KAPA HiFi Real Time Amplification kit and SPRI cleaned again. Our detailed protocols for the above are available on protocols.io^11,12^.

---

**Supplemental Figure 1:**
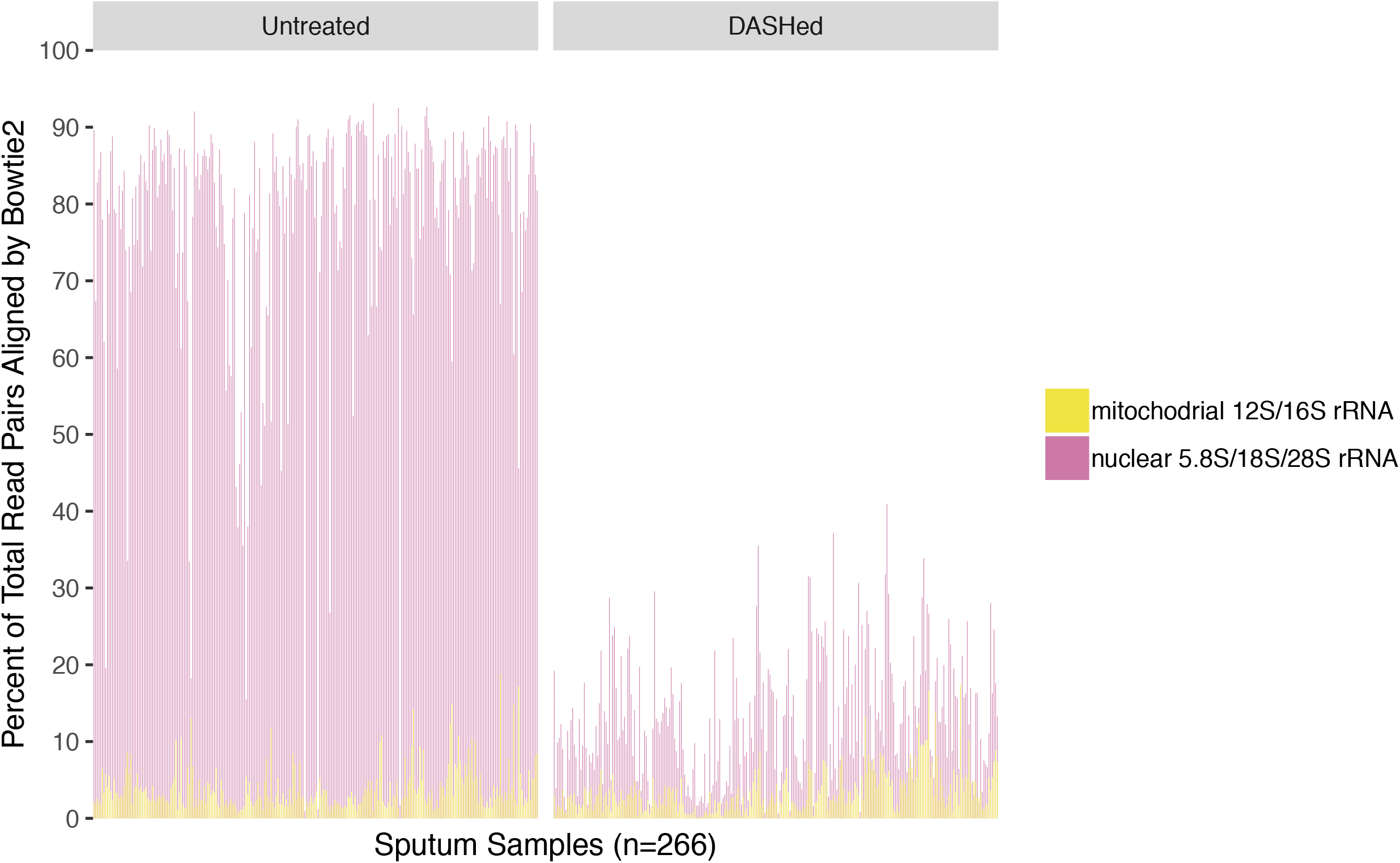
Bowtie2 alignment results to human nuclear ribosomal and mitochondrial RNA for 266 sputum samples in both untreated and DASHed samples. Mitochondrial RNA makes up a small fraction of abundant reads in untreated samples, illustrating a need for a guide set targeting mitochondrial and nuclear ribosomal RNA.

**Supplemental Figure 2:**
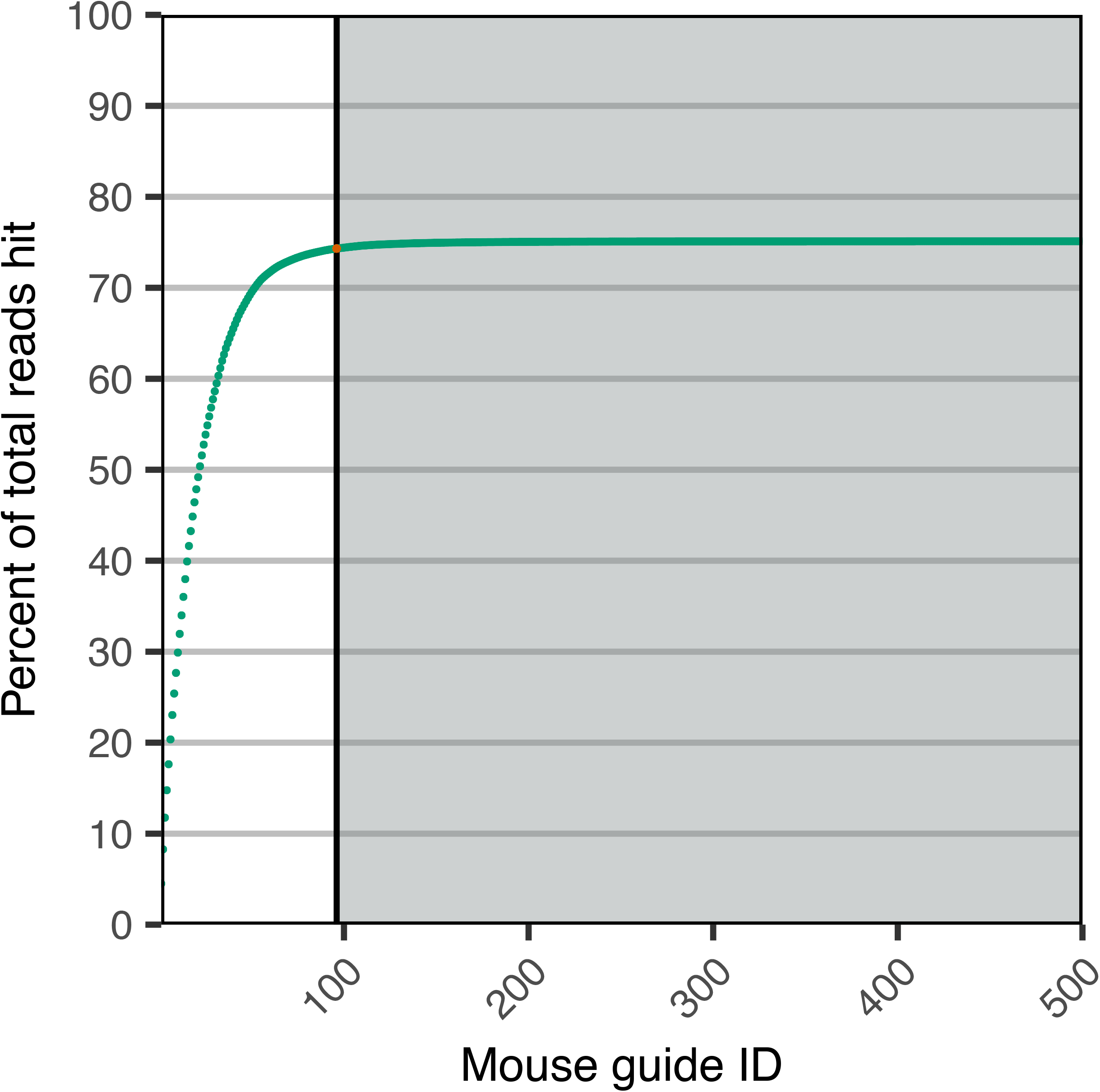
Elbow plot illustrating mouse guides for RNA libraries.

**Supplemental Figure 3:**
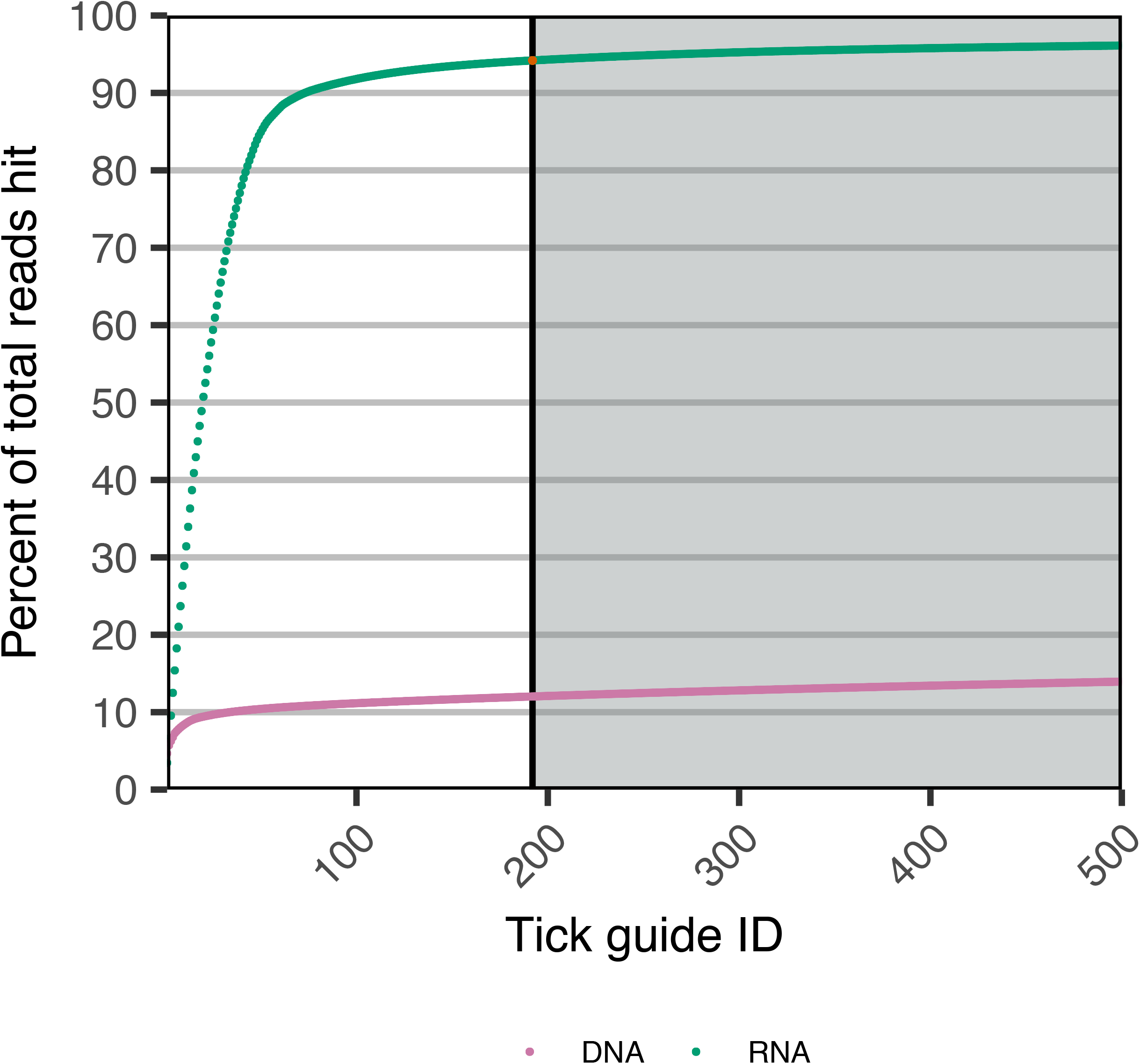
Elbow plot illustrating tick guides for RNA and DNA libraries.

**Table S1:**

| **Virus** | **Sample ID** | **% of reads in untreated targeted by human_ribo gRNAs** | **Virus reads per 10,000 read pairs; untreated** | **Virus reads per 10,000 read pairs; DASHed** | **Fold increase with DASH** |
| --- | --- | --- | --- | --- | --- |
| Betacoronavirus 1 | E083_V6 | 77.04 | 13.60 | 50.77 | 3.73 |
| Human coronavirus 229E | E051_V3 | 91.22 | 20.59 | 379.07 | 18.41 |
|  | E051_V4 | 92.73 | 0.46 | 10.76 | 23.42 |
|  | E051_V5 | 90.33 | 0 | 1.78 | - |
|  | E116_V2 | 91.11 | 6.50 | 60.13 | 9.24 |
| Human respirovirus 3 | E079_V3 | 77.23 | 1.84 | 8.68 | 4.72 |
|  | E079_V4 | 90.15 | 17.54 | 198.02 | 11.29 |
|  | E079_V5 | 86.4 | 1.97 | 6.94 | 3.52 |
| Rhinovirus A | PEX798359 | 82.54 | 1.71 | 5.68 | 3.32 |
|  | E011_V3 | 82.36 | 6.23 | 26.86 | 4.31 |
|  | E011_V4 | 88.98 | 10.65 | 117.29 | 11.01 |
|  | E011_V5 | 89.95 | 1.95 | 25.27 | 12.99 |
|  | E050_V2 | 76.36 | 1.63 | 2.61 | 1.60 |
|  | NB_W19D1 | 74.21 | 16.84 | 141.07 | 8.37 |
|  | E082_V1 | 90.24 | 0 | 7.80 | - |
|  | E100_V1 | 82.14 | 0 | 1.08 | - |
| Rhinovirus B | PEX630531 | 80.81 | 116.36 | 743.79 | 6.39 |
|  | E054_V1 | 86.58 | 1.83 | 14.06 | 7.67 |
| Rhinovirus C | E041_V6 | 79.26 | 6.16 | 65.57 | 10.64 |
|  | E045_V2 | 86.13 | 2.00 | 20.97 | 10.50 |
|  | E063_V2 | 89.88 | 4.20 | 36.77 | 8.76 |
|  | E107_V3 | 88.04 | 0 | 4.28 | - |

## Acknowledgements

We thank Norma Neff and Brian Yu for support with Illumina sequencing; Wei Gu, Eric Chow and Andy May for early work on the DASH method; and Josh Batson for support with software development. We would also like to thank the following researchers involved in preparing the sequencing libraries described in this study: Sputum samples: Samaneh Bolourchi and Suresh Garudadri. Mouse samples: Suzanne Noble and Pauline Basso. Tick samples: Seemay Chou, Anne Sapiro, Valentina Garcia, Amy Kistler and Olga Botvinnik. We thank Dana V. Foss for her helpful comments regarding an earlier draft of this manuscript.

## Protocols.io

IVT reaction: https://www.protocols.io/view/in-vitro-transcription-for-dgrna-3bpgimn

DASH protocol: https://www.protocols.io/view/dash-protocol-6rjhd4n

## Open data

All sequencing data has been deposited in SRA (BioProject PRJNA597373).

All software described can be found here: http://dashit.czbiohub.org

